# CIA: Unveiling Cellular Identities with Cluster-Independent Annotation in Single-Cell RNA Sequencing Data for Comprehensive Cell Type Characterization and Exploration

**DOI:** 10.1101/2023.11.30.569382

**Authors:** Ivan Ferrari, Mattia Battistella, Francesca Vincenti, Andrea Gobbini, Federico Marini, Samuele Notarbartolo, Jole Costanza, Stefano Biffo, Renata Grifantini, Sergio Abrignani, Eugenia Galeota

## Abstract

Single-cell RNA sequencing (scRNA-seq) has revolutionized our understanding of the transcriptional landscape of complex tissues, enabling the discovery of novel cell types and biological functions. However, the identification and classification of cells from scRNA-seq datasets remain significant challenges. To address this, we developed a new computational tool called CIA (Cluster Independent Annotation), which accurately identifies cell types across different datasets without requiring a fully annotated reference dataset or complex machine learning processes. Based on predefined cell type signatures, CIA provides a highly user-friendly and practical solution to functional annotation of single cells. Our results demonstrate that CIA outperforms other state-of-the-art approaches, while also having significantly lower computational running time. Overall, CIA simplifies the process of obtaining reproducible signature-based cell assignments that can be easily interpreted through graphical summaries providing researchers with a powerful tool to explore the complex transcriptional landscape of single cells.

The CIA framework is implemented in both the Python and R programming languages, making it applicable to all main single-cell analysis frameworks, and it is available under the MIT license with its documentation at the following links:

Python package: https://pypi.org/project/cia-python/

Python tutorial: https://cia-python.readthedocs.io/en/latest/tutorial/Cluster_Independent_Annotation.html

R package and tutorial: https://github.com/ingmbioinfo/CIA_R

## Introduction

The emergence of scRNA-seq technology has opened up unprecedented opportunities for exploring gene expression profiles at the individual cell level [1]. scRNA-seq has quickly become the preferred approach for analyzing cell transcriptional readouts, enabling researchers to identify cell transitions, discover new cell types and investigate cellular heterogeneity at various levels of granularity. Understanding cellular diversity is a significant challenge in many areas of biology and biomedical research, including immunology, developmental and stem cell biology, neurobiology, and cancer research [2,3].

Once library preparation and sequencing are completed, count matrices are generated by processing and aligning raw data. These matrices serve as the starting point for a single-cell RNA sequencing analysis. The typical computational workflow involves several key steps: quality control and filtering of cells and genes, normalization of gene expression counts and identification of groups of cells with similar transcriptional patterns [4].

Different clustering methods can be employed to identify these groups, but setting the correct clustering resolution is often challenging [5]. General guidelines may not apply equally well to all datasets [6]. Accurate labeling of cells and clusters is crucial for identifying cell populations, their functions, and the biological states represented in the experimental context. Traditionally, cell type annotation relies on manual curation by domain experts—a process that is time-consuming and may lack reproducibility [7]. This approach depends heavily on prior biological knowledge and is not easily automated. However, efforts are underway to provide annotation guidelines using automated cell labeling tools [8].

To address this challenge, various computational approaches have been developed for automated cell type annotation in scRNA-seq datasets. These methods can be broadly categorized into three main strategies: marker-genes database-based annotation, correlation-based annotation, and supervised classification-based annotation [7]. Marker-genes database-based methods rely on known marker genes and their expression patterns, associated with specific cell types, to assign cell identities. Correlation-based methods compare the gene expression profiles of unlabeled cells or clusters with those of reference datasets to infer cell types based on similarity metrics. Supervised classification-based methods leverage machine learning algorithms trained on labeled reference datasets to classify unlabeled cells [7].

Each strategy offers unique advantages in handling complex datasets and capturing subtle variations in gene expression profiles.

However, the identification of reliable reference datasets or gene signatures is complicated by the high degree of heterogeneity characterizing single-cell datasets, making it challenging to label cell types accurately. This heterogeneity arises from various factors, including technical noise, batch effects, and the intrinsic diversity of cellular states, which can obscure the definition of distinct cell populations [9]. As highlighted by Abdelaal et al. [10], automated annotation methods must account for this variability to improve accuracy and robustness in single-cell analyses. Their study systematically evaluated multiple cell type annotation tools, demonstrating that performance can vary significantly across datasets and highlighting the need for methods that are adaptable to different levels of cellular heterogeneity. This underscores the importance of developing approaches that can accurately classify cells despite dataset-specific variations.

Recently, valuable gene signature collections have been published, covering various organisms, tissues and cell types at different levels of granularity such as ACT [11], CellMarker 2.0 [12], PanglaoDB [13], singleCellBase [14], and CellKb [15]. These collections can be utilized to assign cell types in newly produced single-cell RNA-sequencing datasets. Moreover, consortia are producing large-scale single-cell datasets in the form of atlases to serve as a reliable reference for grasping the underlying differences among cell types, tissues and individuals [16]. The availability of reference atlases, coupled with the ability of machine learning approaches to exploit prior domain knowledge, represents a new way of analyzing single-cell datasets [17].

In this context, we present CIA, a software that automatically assigns accurate cell type labels in scRNA-seq datasets requiring only gene signatures. CIA is a versatile framework that supports comprehensive software packages such as SingleCellExperiment, Seurat and Scanpy, allowing researchers to efficiently annotate cell populations without the need for extensive reference training datasets, thus accelerating the process of biological discovery and interpretation across a wide range of single-cell datasets.

## Results

### The CIA framework and functionality

CIA is a software package developed for the signature-based automatic annotation of cellular populations within scRNA-seq datasets. Designed to operate without relying on clustering, CIA offers a simpler and more streamlined approach to cell classification tools compared to other tools. Among the signature-based methods cited in [7], CIA is the only one focused on annotating single cells, as other methods are cluster-based. Although CellAssign operates at the single-cell level, it still depends on a Bayesian probabilistic model to assign labels to clusters, making it fundamentally cluster-based. In contrast, CIA eliminates the need for clustering, model training, or probabilistic frameworks, providing a direct and efficient tool for high-throughput single-cell RNA-seq data analysis.

We opted for a signature-based approach because it enables quick and efficient exploration of datasets, identifying enrichment for specific signatures of interest without the complexity of training models or needing reference datasets. This makes CIA an excellent tool for rapid analysis, allowing researchers to gain valuable insights without the computational overhead associated with traditional machine learning methods.

CIA takes two main inputs: (i) a dataset with normalized gene expression values for individual cells and (ii) a list of gene signatures with the associated cell types. The software is implemented in both Python and R to accommodate diverse user preferences and frameworks commonly used in the scientific community. Signatures can be supplied in Gene Matrix Transposed (GMT) format from different sources, including URLs, local files, Python dictionaries, or R lists of named vectors. CIA is compatible with major single-cell data formats, including Python AnnData [18], R SingleCellExperiment [19], and SeuratObject [20], and is specifically designed to work with datasets that have been normalized using total count scaling, followed by log-transformation. Tutorials included in the CIA packages, demonstrate the required preprocessing steps. This ensures seamless integration into existing analytical workflows.

CIA is composed of several functions, with the core functions being *score_signature, score_all_signatures* and *CIA_classify*, and other additional functionalities such as performance reporting and classification refinement: (i) *score_signature* calculates scores (*detailed in Methods*) for a given signature for each cell in the dataset, generating a score vector. (ii) *score_all_signatures* calculates scores for multiple signatures returning a matrix of scores. This score matrix, where each entry corresponds to the signature scores for every gene signature and cell in the expression matrix, is crucial for the subsequent classification (**Fig 1**). Notably, the returned output can be effortlessly integrated into the original input object (as continuous sample metadata), reducing the effort to track all operations performed. (*iii*) Exploiting the score matrix, *CIA_classify* labels each single cell of the dataset according to the best matching signature. To prevent misassignment when signatures yield comparable scores, a parameter allows labeling the cell as ‘Unassigned’ if a user-defined similarity threshold is not met (see *Methods*). This approach is critical for maintaining classification specificity and avoiding potential misclassifications, which can lead to erroneous biological conclusions.

**Fig 1.**
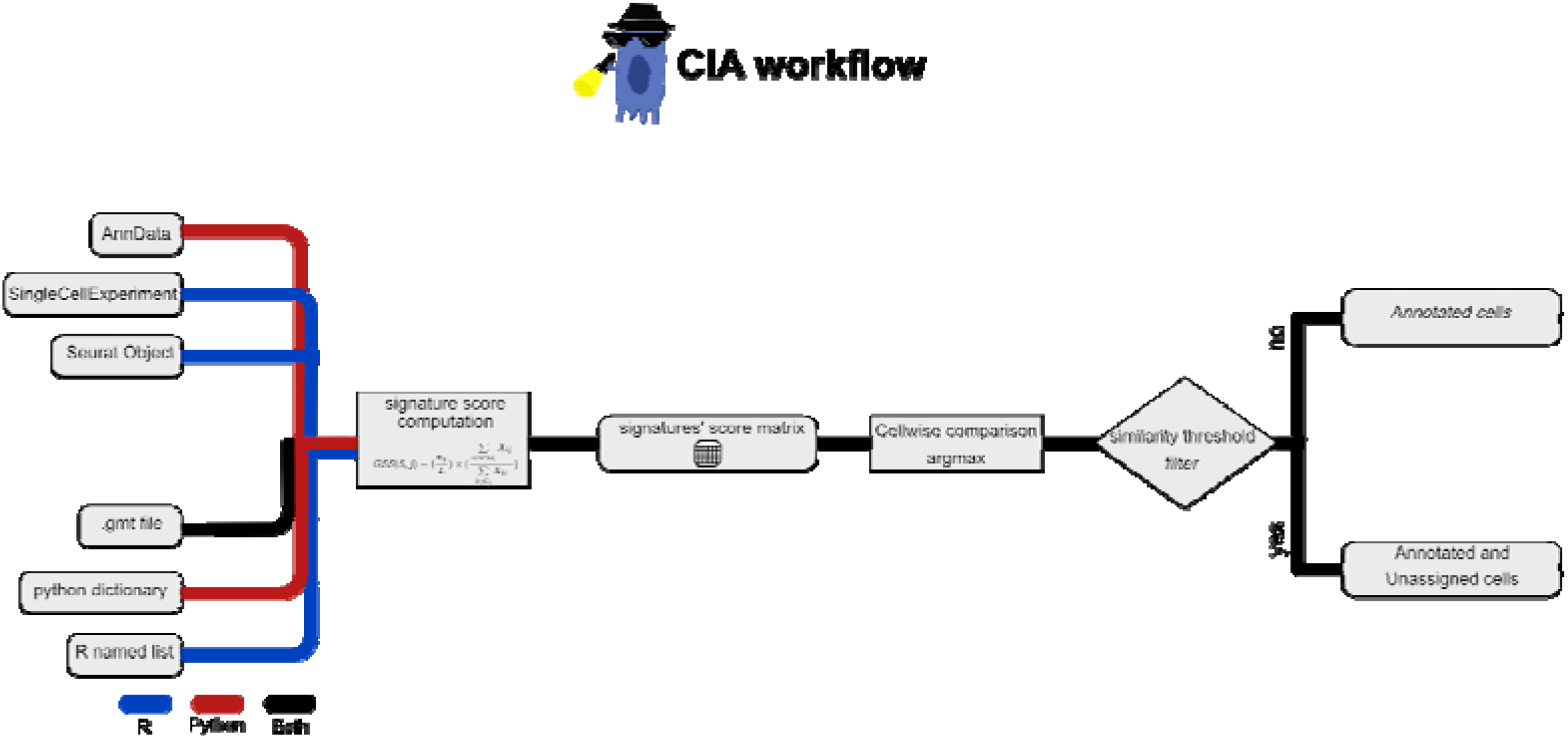
Schematic representation of CIA workflow. CIA requires as inputs a list of named gene signatures and a single-cell object (Anndata, SingleCellExperiment, SeuratObject) containing the normalized gene expression matrix. The signature_score function calculates for each cell and each signature a gene signature score. Then, raw scores from the signature score matrix are scaled in the interval [0, 1]. Finally, CIA_classify assigns cell labels based on the top scored signature name. At user discretion, it is possible to use a similarity threshold to mark as unassigned cells having top scoring values too close.

Moreover, CIA offers additional functionalities to assist users in evaluating both the quality of signatures and the classification performance:

*(i) grouped_distributions* calculates and plots median signature scores in groups of cells (e.g. user-defined annotation) for each signature. The function performs comparative statistical analysis on cell populations using Wilcoxon signed-rank tests and Mann-Whitney U tests to evaluate the discriminative power of gene signatures within and across cell groups. The Wilcoxon signed-rank test checks if each pair of gene signatures has significantly different distributions within the same cell group. The Mann-Whitney U test checks if each gene signature has significantly different distributions across different cell groups. In this way the function identifies potential issues such as inadequate gene signatures or flawed clustering. (*ii*) *group_composition* returns and plots a contingency table of the classification results. It is beneficial when a reference classification is available and can be used to test how labels assigned by CIA align with the reference groups. (*iii*) *classification_metrics* and *grouped_classification_metrics* compute sensitivity, specificity, precision, accuracy and F1-score. *classification_metrics* returns the metrics per cell group, providing information about the classification quality for individual cell populations. *grouped_classification_metrics* offers the same metrics for the entire dataset, allowing for rapid comparison of different cell annotations simultaneously.

Additionally, CIA includes several auxiliary functions: *signatures_similarity, filter_degs*, and *celltypist_majority_vote. signatures_similarity* calculates the similarity between couples of gene signatures using either the Jaccard index or by computing the percentage of overlap of the first signature with the second. *filter_degs* (available only in Python version) subsets differentially expressed genes (DEGs) stored in the “uns” layer of the AnnData object based on different thresholds of log fold changes, z-scores, mean expression and percentage of cells expressing genes. *celltypist_majority_vote* integrates Celltypist [21] code and is optionally used only after CIA workflow. It extends the most represented cell-level annotations to independently defined cell groups. If reference clusters are not provided by the user, the function employs an over-clustering approach using the Leiden algorithm to define groups (see the Celltypist documentation for further details).

### CIA correctly classifies the PBMC3k dataset

We tested CIA on a human PBMC3k single cell dataset from 10X Genomics [22], comprising 2700 cells from a healthy donor. To automatically annotate cells, we utilized differentially expressed genes for specific cell populations, obtained from a publicly available reference atlas of human peripheral mononuclear blood cells (PBMC Atlas) [23], which was annotated through both transcript and protein levels inspections. We chose the PBMC Atlas as the reference for our study due to its comprehensive and accurate annotation of human peripheral blood cell types, which includes a variety of well-characterized immune cell populations at both transcriptomic and proteomic levels. Using such a well-defined atlas provides a solid reference point for cell type classification and allows for a comparative evaluation of CIA’s performance against existing classification methods. Additionally, the PBMC Atlas is widely used in single-cell RNA sequencing (scRNA-seq) research and has a robust dataset that has been integrated with additional information such as cell surface markers, making it particularly suited for training and validating our method.

The differential expression analysis (see Materials and Methods) yielded seven gene signatures, also available in the CIA tutorials and online repository, that represent immune populations: B lymphocytes, CD4+ and CD8+ T lymphocytes, dendritic cells (DC), monocytes (Mono), natural killer cells (NK) and platelets (**Fig 2A**).

**Fig 2.**
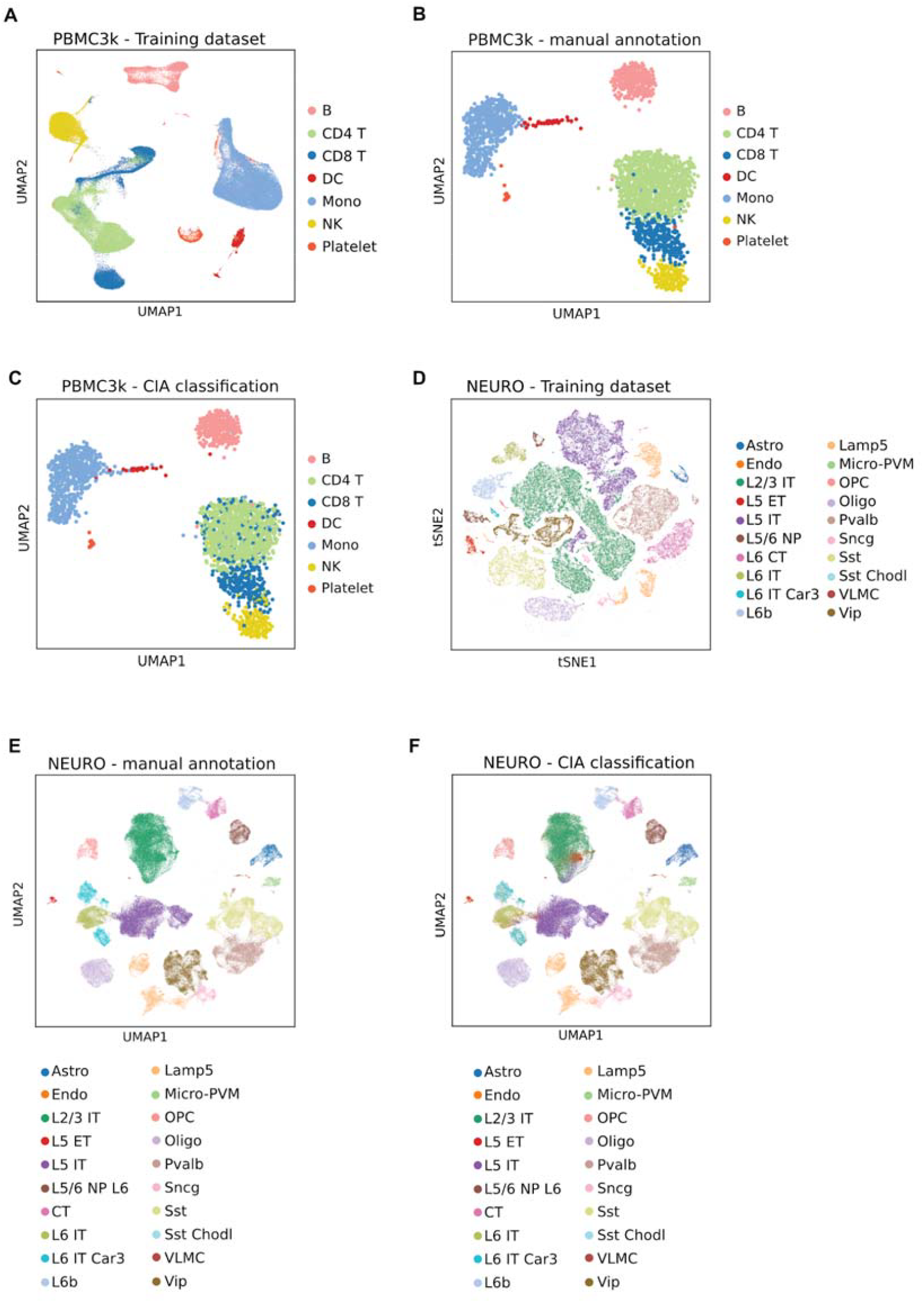
A) UMAP plot showing the PBMC training dataset with the coarser level of annotation. B) UMAP plot showing the PBMC3k dataset with ground truth annotation. C) UMAP plot of PBMC3k dataset showing cell labels predicted by CIA. D) UMAP plot showing the NEURO_training dataset with the coarser level of annotation. E) UMAP plot showing the NEURO_test dataset with ground truth annotation. F) UMAP plot of NEURO_test dataset showing cell labels predicted by CIA.

We benchmarked our results against the manual classification of the PBMC3k dataset (**Fig 2B**). First, we applied the *score_all_signature*s function to calculate scaled scores associated with each cell of the PBMC Atlas for each of the seven subpopulation signatures. By utilizing the *grouped_distributions* we obtained the median signature score for each signature within each population. The function also assessed their statistical significance (**Fig S1A**). We observed that the highest scores for each signature were consistent with the corresponding manually annotated subpopulation.

We then used the signatures obtained from the PBMC Atlas to classify the PBMC3k dataset with *CIA_classify* function (**Fig 2C**). The heatmaps with median signature scores clearly showed that higher median signature scores corresponded to the correct population (**Fig S2A**). As expected, the *grouped_distributions* function returned a warning indicating that the signature score distribution of DC and Mono were not significantly different within the manually annotated dendritic cells cluster. This diagnostic information is valuable because it highlights when signatures lack specificity for the target population. For example, the signature for monocytes had lower median scores due to the lack of specific identity markers, while the DC signature exhibited high variability within the DC cluster.

The statistical tests were crucial for identifying these issues. When a signature is not specific enough, as with the Mono signature, or when scores are highly variable within a population, as with the DC signature, it indicates the need for refinement. To address these problems, it is beneficial to identify more specific population markers or to specify groups of markers that better capture the variability of the given population.

However, since for each distribution, there is only a cell group in which values are significantly higher than all the other groups, the simple inspection of PBMC3k UMAP showing the different signature scores was sufficient to annotate the expected cell populations (**Fig S1B**).

*CIA_classify* inferred cell identities with high accuracy (0.98), precision (0.92) and F1 (0.92) values (**Table 1**) in less than 0.30 seconds with different computational setups.

**Table 1.** Table reporting cell-level classification performances. * Running time of differential expression analysis (23.3s) and DEGs filtering (20.3s) are not included since signature refinement is not the intent of this package and it should be done before CIA classification.

|  | Classifier | Classification performance |  |  |  |  | Running time (s) |  |  |  |  |  |  |
| --- | --- | --- | --- | --- | --- | --- | --- | --- | --- | --- | --- | --- | --- |
|  |  | SE | SP | PR | ACC | F1 | 1 CPUs | 2 CPUs | 4 CPUs | 8 CPUs | 16 CPUs | 32 CPUs | 64 CPUs |
| PBMC3k | CIA_Python | 0.92 | 0.99 | 0.92 | 0.96 | 0.92 | 0.28 | 0.25 | 0.21 | 0.20 | 0.20 | 0.20 | 0.20 |
|  | CIA_R | 0.92 | 0.99 | 0.92 | 0.96 | 0.92 | 2.14 | 2.15 | 2.06 | 2.02 | 2.01 | 2.00 | 1.99 |
|  | SingleR | 0.76 | 0.96 | 0.77 | 0.93 | 0.76 | 390.14 | 294.92 | 219.98 | 195.31 | 158.75 | 144.66 | 138.17 |
|  | Garnett | 0.24 | 0.94 | 0.39 | 0.84 | 0.29 | 1365.96 | 1258.36 | 955.40 | 820.24 | 908.12 | 827.89 | 667.59 |
|  | Celltypist | 0.75 | 0.96 | 0.75 | 0.93 | 0.75 | 2759.78 | 2784.00 | 1811.60 | 1412.93 | 1200.72 | 1202.65 | 1156.06 |
|  | AUCell | 0.65 | 0.94 | 0.65 | 0.90 | 0.65 | 4.94 | 4.56 | 3.65 | 3.50 | 3.50 | 3.41 | 3.40 |
| NEURO | CIA_Python | 0.93 | 1.00 | 0.93 | 0.99 | 0.93 | 1.62 | 1.72 | 1.13 | 0.83 | 0.63 | 0.74 | 0.51 |
|  | CIA_R | 0.93 | 1.00 | 0.93 | 0.99 | 0.93 | 64.72 | 60.26 | 41.70 | 33.37 | 29.53 | 27.04 | 25.29 |
|  | SingleR | 0.97 | 1.00 | 0.99 | 1.00 | 0.98 | 7990.20 | 5041.95 | 2941.85 | 2558.36 | 1438.06 | 925.64 | 814.31 |
|  | Garnett | 0.03 | 0.98 | 0.09 | 0.98 | 0.06 | 6768.94 | 5679.33 | 4102.17 | 3639.62 | 3519.80 | 3546.61 | 3523.84 |
|  | Celltypist | 0.85 | 0.99 | 0.85 | 0.98 | 0.85 | 13542.59 | 13261.55 | 7071.65 | 3798.50 | 2398.14 | 2219.50 | 1364.89 |
|  | AUCell | 0.86 | 0.99 | 0.86 | 0.99 | 0.86 | 469.71 | 469.91 | 280.66 | 236.61 | 239.05 | 229.18 | 235.30 |

### Comparison of CIA with state-of-art classifiers

We compared the performance of CIA with two machine learning-based algorithms, Celltypist [21] and Garnett [24], as well as a reference-based classifier, SingleR [25], on two test datasets: PBMC3k and NEURO_test [26], a dataset of 119,152 nuclei from Middle Temporal Gyrus (MTG) derived from 5 post-mortem human brains (see *Methods)* (**Table1, Fig. 2D-F, Figure S3**). The NEURO_test dataset is chosen in order to assess the CIA capability to handle larger and newly generated data from big consortia, with more biologically complexity and more granular annotations. Additionally, we evaluated AUCell [27], a popular signature scoring method, by assigning each cell the label of the signature with the highest AUCell score. However, AUCell scores are not directly comparable across different signatures, even within the same dataset, due to their dependency on gene set size and expression distribution. Larger gene sets tend to yield higher scores, while smaller or lower-expressed signatures may receive lower scores, making it difficult to determine the best-matching signature for a given cell. As a result, AUCell requires the manual selection of signature-specific thresholds, preventing a fully automated and standardized annotation process. Moreover, since multiple signatures can have high scores for the same cell, AUCell does not inherently provide a unique classification output. In contrast, CIA directly assigns a unique cell type label by identifying the most compatible signature, ensuring comparability across signatures and offering a principled way to handle unassigned cells when signatures are ambiguous.

For Celltypist, Garnett, and SingleR, we used the PBMC Atlas as a training dataset to classify PBMC3k and a dataset of 76,533 human primary motor cortex nuclei from the Allen Institute for Brain Science [28] (Neuro_training, Fig 2D) to classify NEURO_test. Signatures obtained from the PBMC Atlas (https://github.com/ingmbioinfo/cia/blob/master/docs/tutorial/data/atlas.gmt) were used by CIA and AUCell to annotate cells in the PBMC3k dataset, while signatures for classifying the Neuro_test dataset were retrieved directly from the Azimuth web portal by selecting subclass annotations of the human motor cortex dataset [29] and saving them to a GMT file. Comparing the performances of all the classifiers in the labeling of the PBMC3k dataset, CIA clearly outperformed all the others both in terms of classification accuracy and running time. Concerning the NEURO_test, CIA inferred cell identities with high accuracy (0.99), precision (0.93) and F1 (0.93) values, showing its reliability when using publicly available signatures. CIA F1 value stands only 0.05 points below SingleR, that reaches a nearly perfect cell labeling (F1=0.98). In this classification, CIA outperformed all the other classifiers in terms of running time, being able to classify around 120k cells in 1 minute (less than 1 second using CIA_python with 8 CPUs). Notably, Garnett exhibited the poorest classification performance with a runtime comparable to SingleR. We hypothesized that the underlying algorithm struggles with managing large signatures, and modifications to the specified parameters only slightly improved performances. Taken together, all these results show the consistent highly accurate classification of CIA (both classification tasks completed with F1 > 0.91, while the second top scoring SingleR F1=0.76 and F1=0.98) and a significantly faster labeling compared to all other methods.

### CIA efficiently highlights unknown cell types

To further test the ability of our classifier to detect the presence of unanticipated cell types, we performed a classification on the Seurat human cord blood mononuclear cells (CBMC) from the “Using Seurat with Multimodal data” vignette (https://satijalab.org/seurat/articles/multimodal_vignette) (**Fig 3A-B**).

**Fig 3.**
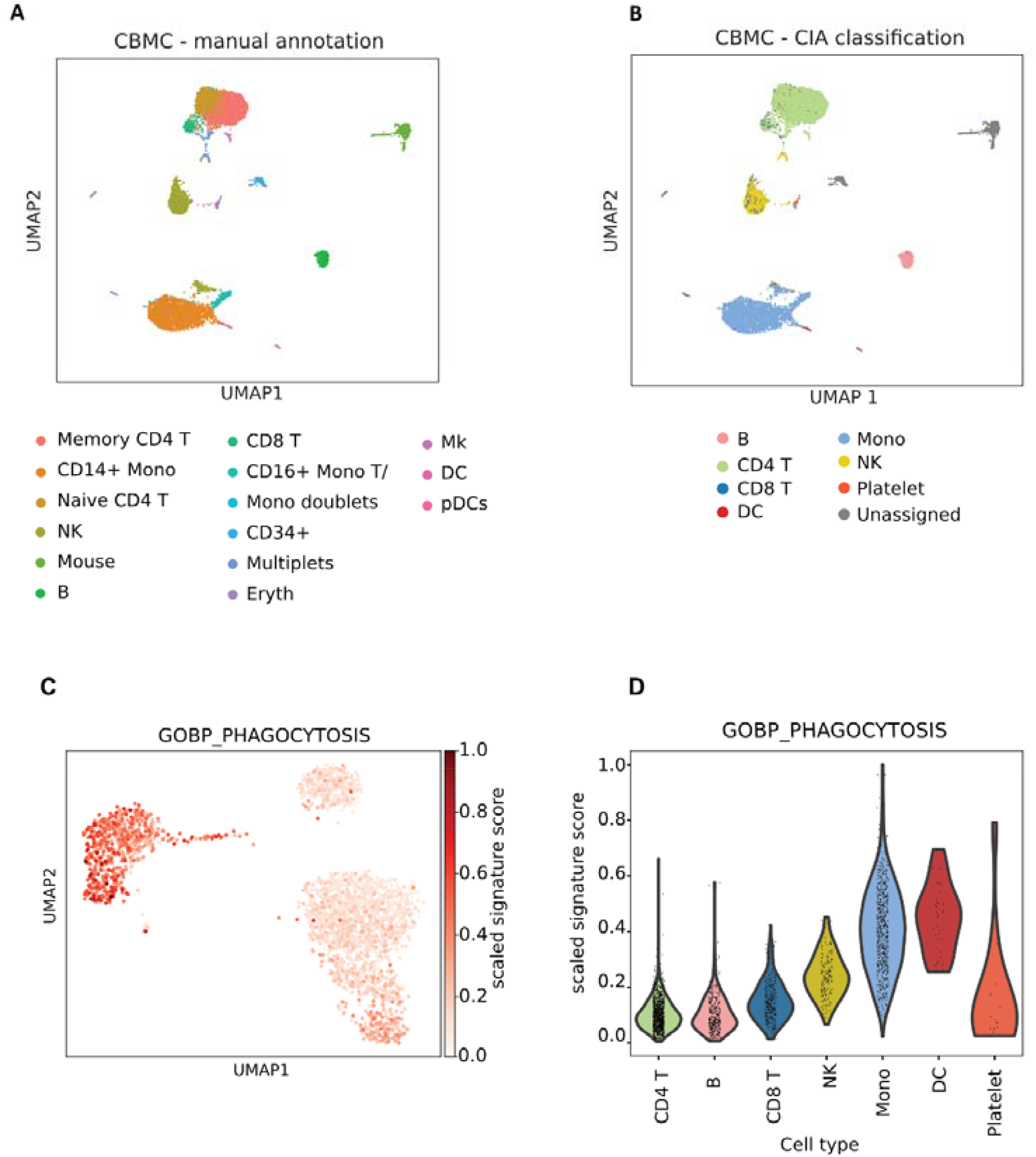
A) UMAP plot showing the CBMC dataset with the ground truth annotation. B) UMAP plot of CBMC dataset showing cell labels predicted by CIA using PBMC atlas signature. Similarity threshold labels as unassigned (gray) Mouse cells and doublets. C) UMAP plot of PBMC3k showing the scaled signature scores of “GOBP_PHAGOCYTOSIS”. D) Violin plot of the scaled signature score distributions of “GOBP_PHAGOCYTOSIS” grouped by ground truth labels.

The CBMC dataset contains human blood cell types and additional mouse cells (∼5%) used as negative controls. Signatures previously extracted from the PBMC Atlas were used for the classification. Although the PBMC Atlas signatures represent cell populations at lower granularity compared to the annotations provided with the CBMC dataset, CIA was able to correctly distinguish such populations. Strikingly, we correctly labeled as “Unassigned” those cells whose transcriptional profiles were not represented by the provided signatures (78.45% of mouse cells and 90.38% of human erythrocytes).

### CIA enables the fast investigation of functional signatures

Beyond cell type recognition across datasets, we explored the capability of CIA to identify cells exhibiting specific biological functions. To this end, we calculated the scaled signature score of the “GOBP_PHAGOCYTOSIS” (GO:0006909) signature from the Molecular Signature Database [28] (Fig 3C). By providing the URL pointing to the GMT file to the score_signature function, we obtained a quantitative layer representing phagocytic activity, which was then superimposed on the UMAP visualization of the entire dataset.

The highest score values were observed in the clusters corresponding to Monocytes (Mono) and Dendritic Cells (DC), which are known to exhibit phagocytic activity (Mann-Whitney U test p<0.01), while all the other cell groups had low scores (Fig 3D). These results demonstrate how CIA can be effectively used for rapid functional characterization at single cell resolution.

## Methods

### Signature score

The *score_all_signatures* function calculates signature scores as previously described in [29]. In brief, this approach takes as input an object with raw gene counts matrix as AnnData, SeuratObject or SingleCellExperiment object, a dictionary of gene signatures (either directly or in GMT format) and an option to set the score modality. The function calls *score_signature*, in which each cell j is evaluated to determine how representative it is of the predefined gene signature S. Our scoring approach involves the following steps:

First we calculate the proportion of genes within the signature S that are expressed in cell j.

Next, we determine the relative contribution of the expressed signature genes to the total gene expression in cell j. Specifically we sum expression counts of the genes in S that are expressed in cell j and divide this sum by the total expression count of all genes expressed in cell j.

The final score thus reflects both the proportion of signature genes that are expressed and their relative contribution to the total gene expression within the cell.

The “raw” gene signature score (GSS) for signature S and cell j is defined as:

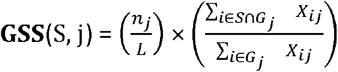

Where:

⍰ S is the set of genes in the signature;

⍰ n_j_ is the number of genes in the signature S that have non-zero expression in cell *j*;

⍰ L is the total number of genes in the signature S;

⍰ *G*_*j*_ is the set of genes expressed (with counts > 0) in cell *j*;

⍰ X is the gene expression matrix where *X*_*ij*_ represents the expression of gene i in cell j.

The score mode input can be used to specify the type of scores to be computed, with options such as “raw”, “scaled”,”log”, “log2” or “log10”.

When the option score_mode is set to “scaled”, raw scores of each cell are divided by the maximum raw score in the dataset, in order to obtain values in the range [0-1].

### Classification

The *CIA_classify* function uses *score_all_signatures* in *scaled* mode to compute the scores for input gene signatures and subsequently classify each cell based on these scores. Each cell is assigned the label of the signature with the highest score. To enhance classification accuracy, the function includes a *similarity_threshold* parameter. This threshold specifies the minimum difference required between the highest and the second-highest scores for a confident assignment. If the difference is below this threshold, the cell is labeled as ‘Unassigned’. The function also supports parallel processing through the *n_cpus* parameter, enabling efficient computation on large datasets.

### Score distribution analysis

The function *grouped_distributions* can be used to determine if the distributions of scores associated with a particular signature are significantly different from others. For each annotated group of cells (e.g. clusters), considered as ground truth annotations, this function applies a Wilcoxon test to verify if the distribution of the scores associated to the signature with the highest median is significantly different from the distributions of scores associated with other signatures. The function then tests, by Mann-Whitney U test, if the distribution of scores of the best signature for each cluster is significantly different from the distribution of scores for the same signature in other clusters. The function returns a heatmap plot of the median scores for each signature in each group, and a warning message if one of the tests has a p-value > 0.01.

### Data processing

#### Extraction of signatures from PBMC Atlas

Cell population signatures for the classification of PBMC3k dataset were extracted from the PBMC Atlas [23], a single-cell atlas including 161764 cells. We omitted 6789 cells labeled as ‘other T’ since they are not present in the PBMC3k dataset. The ‘other’ cluster, predominantly platelets, was relabeled as ‘Platelet’ for clarity. Based on the recent weighted nearest neighbor framework, this atlas integrates gene expression measurements with those of cell surface proteins quantified by the feature barcoding (10x technology), thus providing a very specific and reliable annotation. Cell populations of the atlas include three different levels of granularity. For our analyses, the coarser level was chosen (level I). To extract signatures from the atlas we performed a classical differential expression analysis (DEA) exploiting the *scanpy*.*tl*.*rank*.*genes_groups* (with default parameters) of the remaining annotated populations (B lymphocytes, CD4+ and CD8+ T lymphocytes, dendritic cells, monocytes, natural killer cells and “other” cells, mainly composed by platelets). DEG lists were subsequently filtered through the *filter_degs* to set thresholds on the fold-change (1.5), expression mean (0.25), z-score (5) and percentage of cells within each cluster which express the gene (40%).

#### NEURO_test dataset preparation

NEURO_test dataset [24] was built starting from the AnnData Object “Human MTG 10x SEA-AD” dataset available on the Allen Brain Map website (https://portal.brain-map.org/). Since the Azimuth signatures are not meant to distinguish “Chandelier” or “Pax6+” progenitor cells from the others, those cells have been removed to avoid the misinterpretation of results.

#### Evaluation of CIA performances and comparison with other classifiers

For classification performance evaluation, we employed the *classification_metrics* function to calculate sensitivity, specificity, precision, accuracy, and F1-scores. These metrics were derived by comparing the classifications from CIA, Celltypist (v1.6.2), Garnett (v0.2.20), SingleR(v2.4.1) and AUCell (implemented in the Python package OmicVerse [31] v1.5.6) with the ground truth annotations provided by the datasets. To assess the computational efficiency, we measured the running time of the classification process while incrementing the number of CPUs utilized (1, 2, 4, 8, 16, 32, 64) on a single high performance computing node equipped with 256 GB of RAM. Each configuration was tested 10 times to account for run-to-run variability and ensure reliability of the results. Celltypist classifies cells by applying a combined approach based on logistic regression and stochastic gradient descent. The Celltypist model was trained using the *celltypist*.*train* function with feature selection enabled. Subsequently, the model was applied to the test dataset using the *celltypist*.*annotate* function. Garnett trains a classifier using elastic net multinomial regression. Models were trained on the PBMC atlas and a NEURO_training dataset using the *train_cell_classifier* function with *min_observations* set to 50 and *max_training_samples* set to 800. Cell assignment on the PBMC3k and the NEURO_test dataset was performed using the *classify_cells* function. Concerning the SingleR package, the *SingleR* function, which is a wrapper of *trainSingleR* and *classifySingleR* functions, was applied with default parameters. AUCell was evaluated employing the function *ov*.*single*.*pathway_aucell* to obtain AUC scores. Once AUCell scores were obtained, cells were classified according to the top-scoring signature.

## Discussion

In this study we developed a novel computational tool called CIA, which accurately identifies cell types across diverse datasets relying on a fully annotated reference dataset or complex machine learning processes. Our approach lays its foundation on the unique transcriptional signatures associated with each cell type, enabling CIA to assign cell type labels with high accuracy by comparing the expression profiles of query cells against a set of pre-computed signatures.

CIA is designed for ease of use, featuring comprehensive documentation that facilitates its integration into existing single-cell analysis workflows. Its open-source code is freely available on GitHub, with implementations in both Python (https://github.com/ingmbioinfo/cia) and R (https://github.com/ingmbioinfo/CIA_R). This dual compatibility allows for seamless use alongside popular single-cell analysis tools such as Scanpy and Seurat. One of the distinguishing features of CIA is its independence from the clustering step, empowering users to explore biologically relevant cell groups prior to defining clustering resolutions. The rapid classification capabilities of CIA enable it to handle large datasets in seconds, providing an additional layer of prior knowledge that clustering algorithms often overlook. This capability allows for more confident refinement of cluster boundaries, ultimately enhancing the understanding of the dataset under investigation.

While pretrained models like scBERT [32] and scGPT [33] are powerful tools in single-cell research, CIA focuses on simplicity, accessibility, and transparency. It avoids the ‘black-box’ nature of deep learning by relying on interpretable gene signatures and an intuitive algorithm. CIA is lightweight, requiring only standard computational resources, and is particularly suited for researchers prioritizing rapid and resource-efficient classification without extensive training datasets. By complementing rather than competing with deep learning models, CIA offers a streamlined, user-friendly solution for cell-type annotation.

Unlike many existing tools, CIA does not require a training dataset derived from similar conditions, which can introduce bias associated with specific datasets. By mitigating these issues, CIA facilitates accurate and efficient cell labeling. A notable aspect of CIA is the inclusion of a similarity threshold that can prevent the classification of cells with scores that are closely aligned among signatures. This feature is particularly advantageous as it highlights cells for which the signatures may be inappropriate or those exhibiting ambiguous transcriptional profiles. Consequently, it helps to avert misclassification and uncovers potential new cell types or biological conditions that might otherwise be overlooked.

As a proof of concept, we demonstrated the efficacy of our classifier using the CBMC dataset annotation, showing that CIA can accurately detect cell populations corresponding to the input signatures, while designating those that do not fit as “Unassigned” negative controls. Overall, our method outperformed other state-of-the-art approaches while significantly reducing computational runtime and enhancing interpretability. Although the classification of “Unassigned” cells may appear as a reduction in performance, it offers an opportunity to generate new hypotheses regarding the nature of these cells within single-cell datasets. Given the proliferation of signature databases and large community cell atlases, computational efficiency may become a critical consideration. The straightforward workflow, coupled with rapid execution and the interpretability afforded by gene signatures, positions CIA as a valuable asset in the era of big omics.

In addition to its core modules, CIA provides several utilities that enable users to visualize signature enrichment scores on UMAPs, generate contingency matrices, compare classifications against reference datasets, evaluate classification performance, and plot classification results on UMAPs. By integrating this information into existing AnnData, Seurat, or SingleCellExperiment objects, CIA simplifies the graphical representation of signature enrichment scores and classification outcomes, thereby streamlining incorporation into any of the established single-cell workflows. Furthermore, in addition to being a robust alternative to cluster-based methods, CIA can assist in annotating independently obtained clusters. By natively integrating the majority vote feature from Celltypist [21], it also harmonizes labels within clusters, ensuring consistent and accurate annotations. Using standardized types to store data and the computed results, the output of CIA can be seamlessly further explored in interactive applications such as iSEE [34] or CELLxGENE [35], as demonstrated in the package vignette.

Given the ease with which signature-shaped information can be generated or retrieved within the current and future single-cell landscape [17], we anticipate that CIA will become increasingly valuable for obtaining quantitative overviews, ultimately enhancing the extraction of insights and comprehensive interpretation of data. Our findings suggest that CIA is an effective tool for scRNA-seq analysis, enabling researchers to identify cell types and biological functions in their datasets without the need for extensive training or optimization, all within a relatively short time frame. Additionally, we expect CIA’s performance to scale linearly with increasing cell numbers, making it amenable for use on standard laptops equipped with typical computational resources.

## References

1. Chen G, Ning B, Shi T. Single-Cell RNA-Seq Technologies and Related Computational Data Analysis. Front Genet. 2019;10: 317. doi:10.3389/fgene.2019.00317

2. Nguyen QH, Pervolarakis N, Nee K, Kessenbrock K. Experimental Considerations for Single-Cell RNA Sequencing Approaches. Front Cell Dev Biol. 2018;6: 108. doi:10.3389/fcell.2018.00108

3. Wagner A, Regev A, Yosef N. Revealing the vectors of cellular identity with single-cell genomics. Nat Biotechnol. 2016;34: 1145–1160. doi:10.1038/nbt.3711

4. Amezquita RA, Lun ATL, Becht E, Carey VJ, Carpp LN, Geistlinger L, et al. Orchestrating single-cell analysis with Bioconductor. Nat Methods. 2020;17: 137–145. doi:10.1038/s41592-019-0654-xù

5. Grabski IN, Street K, Irizarry RA. Significance analysis for clustering with single-cell RNA-sequencing data. Nat Methods. 2023;20: 1196–1202. doi:10.1038/s41592-023-01933-9

6. Duò A, Robinson MD, Soneson C. A systematic performance evaluation of clustering methods for single-cell RNA-seq data. F1000Res. 2020;7: 1141. doi:10.12688/f1000research.15666.3

7. Pasquini G, Rojo Arias JE, Schäfer P, Busskamp V. Automated methods for cell type annotation on scRNA-seq data. Computational and Structural Biotechnology Journal. 2021;19: 961–969. doi:10.1016/j.csbj.2021.01.015

8. Clarke ZA, Andrews TS, Atif J, Pouyabahar D, Innes BT, MacParland SA, et al. Tutorial: guidelines for annotating single-cell transcriptomic maps using automated and manual methods. Nat Protoc. 2021;16: 2749–2764. doi:10.1038/s41596-021-00534-0

9. Choi YH, Kim JK. Dissecting Cellular Heterogeneity Using Single-Cell RNA Sequencing. Mol Cells. 2019;42: 189–199. doi:10.14348/molcells.2019.2446

10. Abdelaal T, Michielsen L, Cats D, Hoogduin D, Mei H, Reinders MJT, et al. A comparison of automatic cell identification methods for single-cell RNA sequencing data. Genome Biol. 2019;20: 194. doi:10.1186/s13059-019-1795-z

11. Quan F, Liang X, Cheng M, Yang H, Liu K, He S, et al. Annotation of cell types (ACT): a convenient web server for cell type annotation. Genome Med. 2023;15: 91. doi:10.1186/s13073-023-01249-5

12. Hu C, Li T, Xu Y, Zhang X, Li F, Bai J, et al. CellMarker 2.0: an updated database of manually curated cell markers in human/mouse and web tools based on scRNA-seq data. Nucleic Acids Research. 2023;51: D870–D876. doi:10.1093/nar/gkac947

13. Franzén O, Gan L-M, Björkegren JLM. PanglaoDB: a web server for exploration of mouse and human single-cell RNA sequencing data. Database. 2019;2019. doi:10.1093/database/baz046

14. Meng F-L, Huang X-L, Qin W-Y, Liu K-B, Wang Y, Li M, et al. singleCellBase: a high-quality manually curated database of cell markers for single cell annotation across multiple species. Biomark Res. 2023;11: 83. doi:10.1186/s40364-023-00523-3

15. Patil A, Patil A. CellKb Immune: a manually curated database of mammalian hematopoietic marker gene sets for rapid cell type identification. 2020. doi:10.1101/2020.12.01.389890

16. Lotfollahi M, Naghipourfar M, Luecken MD, Khajavi M, Büttner M, Wagenstetter M, et al. Mapping single-cell data to reference atlases by transfer learning. Nat Biotechnol. 2022;40: 121–130. doi:10.1038/s41587-021-01001-7

17. Hemberg M, Marini F, Ghazanfar S, Ajami AA, Abassi N, Anchang B, et al. Insights, opportunities and challenges provided by large cell atlases. arXiv; 2024. doi:10.48550/ARXIV.2408.06563

18. Wolf FA, Angerer P, Theis FJ. SCANPY: large-scale single-cell gene expression data analysis. Genome Biol. 2018;19: 15. doi:10.1186/s13059-017-1382-0

19. Aaron Lun [Aut C. SingleCellExperiment. Bioconductor; 2017. doi:10.18129/B9.BIOC.SINGLECELLEXPERIMENT

20. Butler A, Hoffman P, Smibert P, Papalexi E, Satija R. Integrating single-cell transcriptomic data across different conditions, technologies, and species. Nat Biotechnol. 2018;36: 411–420. doi:10.1038/nbt.4096

21. Domínguez Conde C, Xu C, Jarvis LB, Rainbow DB, Wells SB, Gomes T, et al. Cross-tissue immune cell analysis reveals tissue-specific features in humans. Science. 2022;376: eabl5197. doi:10.1126/science.abl5197

22. Zheng GXY, Terry JM, Belgrader P, Ryvkin P, Bent ZW, Wilson R, et al. Massively parallel digital transcriptional profiling of single cells. Nat Commun. 2017;8: 14049. doi:10.1038/ncomms14049

23. Hao Y, Hao S, Andersen-Nissen E, Mauck WM, Zheng S, Butler A, et al. Integrated analysis of multimodal single-cell data. Cell. 2021;184: 3573-3587.e29. doi:10.1016/j.cell.2021.04.048

24. Pliner HA, Shendure J, Trapnell C. Supervised classification enables rapid annotation of cell atlases. Nat Methods. 2019;16: 983–986. doi:10.1038/s41592-019-0535-3

25. Aran D, Looney AP, Liu L, Wu E, Fong V, Hsu A, et al. Reference-based analysis of lung single-cell sequencing reveals a transitional profibrotic macrophage. Nat Immunol. 2019;20: 163–172. doi:10.1038/s41590-018-0276-y

26. Gabitto M, Travaglini K, Ariza J, Kaplan E, Long B, Rachleff V, et al. Integrated multimodal cell atlas of Alzheimer’s disease. 2023. doi:10.21203/rs.3.rs-2921860/v1

27. Aibar S, González-Blas CB, Moerman T, Huynh-Thu VA, Imrichova H, Hulselmans G, et al. SCENIC: single-cell regulatory network inference and clustering. Nat Methods. 2017;14: 1083–1086. doi:10.1038/nmeth.4463

28. Bakken TE, Jorstad NL, Hu Q, Lake BB, Tian W, Kalmbach BE, et al. Comparative cellular analysis of motor cortex in human, marmoset and mouse. Nature. 2021;598: 111–119. doi:10.1038/s41586-021-03465-8

29. Liberzon A, Subramanian A, Pinchback R, Thorvaldsdóttir H, Tamayo P, Mesirov JP. Molecular signatures database (MSigDB) 3.0. Bioinformatics. 2011;27: 1739–1740. doi:10.1093/bioinformatics/btr260

30. Della Chiara G, Gervasoni F, Fakiola M, Godano C, D’Oria C, Azzolin L, et al. Epigenomic landscape of human colorectal cancer unveils an aberrant core of pan-cancer enhancers orchestrated by YAP/TAZ. Nat Commun. 2021;12: 2340. doi:10.1038/s41467-021-22544-y

31. Zeng Z, Ma Y, Hu L, Tan B, Liu P, Wang Y, et al. OmicVerse: a framework for bridging and deepening insights across bulk and single-cell sequencing. Nat Commun. 2024;15: 5983. doi:10.1038/s41467-024-50194-3

32. Yang F, Wang W, Wang F, Fang Y, Tang D, Huang J, et al. scBERT as a large-scale pretrained deep language model for cell type annotation of single-cell RNA-seq data. Nat Mach Intell. 2022;4: 852–866. doi:10.1038/s42256-022-00534-z

33. Cui H, Wang C, Maan H, Pang K, Luo F, Duan N, et al. scGPT: toward building a foundation model for single-cell multi-omics using generative AI. Nat Methods. 2024;21: 1470–1480. doi:10.1038/s41592-024-02201-0

34. Rue-Albrecht K, Marini F, Soneson C, Lun ATL. iSEE: Interactive SummarizedExperiment Explorer. F1000Res. 2018;7: 741. doi:10.12688/f1000research.14966.1

35. Megill C, Martin B, Weaver C, Bell S, Prins L, Badajoz S, et al. cellxgene: a performant, scalable exploration platform for high dimensional sparse matrices. 2021. doi:10.1101/2021.04.05.438318

