## Supplementary Figures for "CIA: Unveiling Cellular Identities with Cluster-Independent Annotation in Single-Cell RNA Sequencing Data for Comprehensive Cell Type Characterization and Exploration"

**Over-clustering and majority voting improve cell-level annotation**

Besides single cell-level annotation, Celltypist can refine cell identities within local subclusters after an over-clustering step. For each subcluster, the label of the most abundant cell type is extended to the whole cell group. Since this additional step might improve the classification performances, we decided to provide the majority voting strategy by wrapping Celltypist function. Subsequently we compared our performances with those of Besca and Celltypist. In detail, over-clustering is performed by the Leiden algorithm, whose resolution is automatically set depending on the number of cells in the dataset.

With this approach, PBMC3k dataset has been divided into 68 small clusters composed of highly similar cells. This new layer of information was then used to ameliorate the previous cell-level classifications of each tool, assigning the most abundant cell type label to each cluster.

Notably, the performances significantly improved for all the tested tools (Table S1). In particular, CIA performed the best classifications, followed by Besca. These results suggest that when clustering is used to refine an already accurate automatic cell-level labelling, the annotation process is facilitated and less sensitive to arbitrary decisions on the cell populations’ boundaries. In this way the knowledge provided by widely established population markers can be complemented by the information of overall transcriptional similarity that the clustering provides.

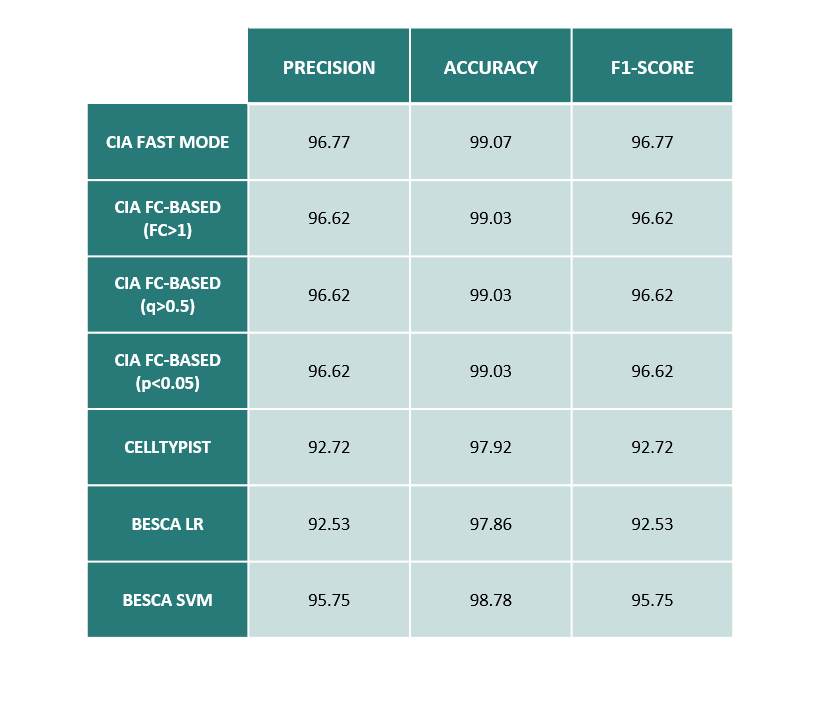

**Table S1.** Table reporting classification performances with majority voting.

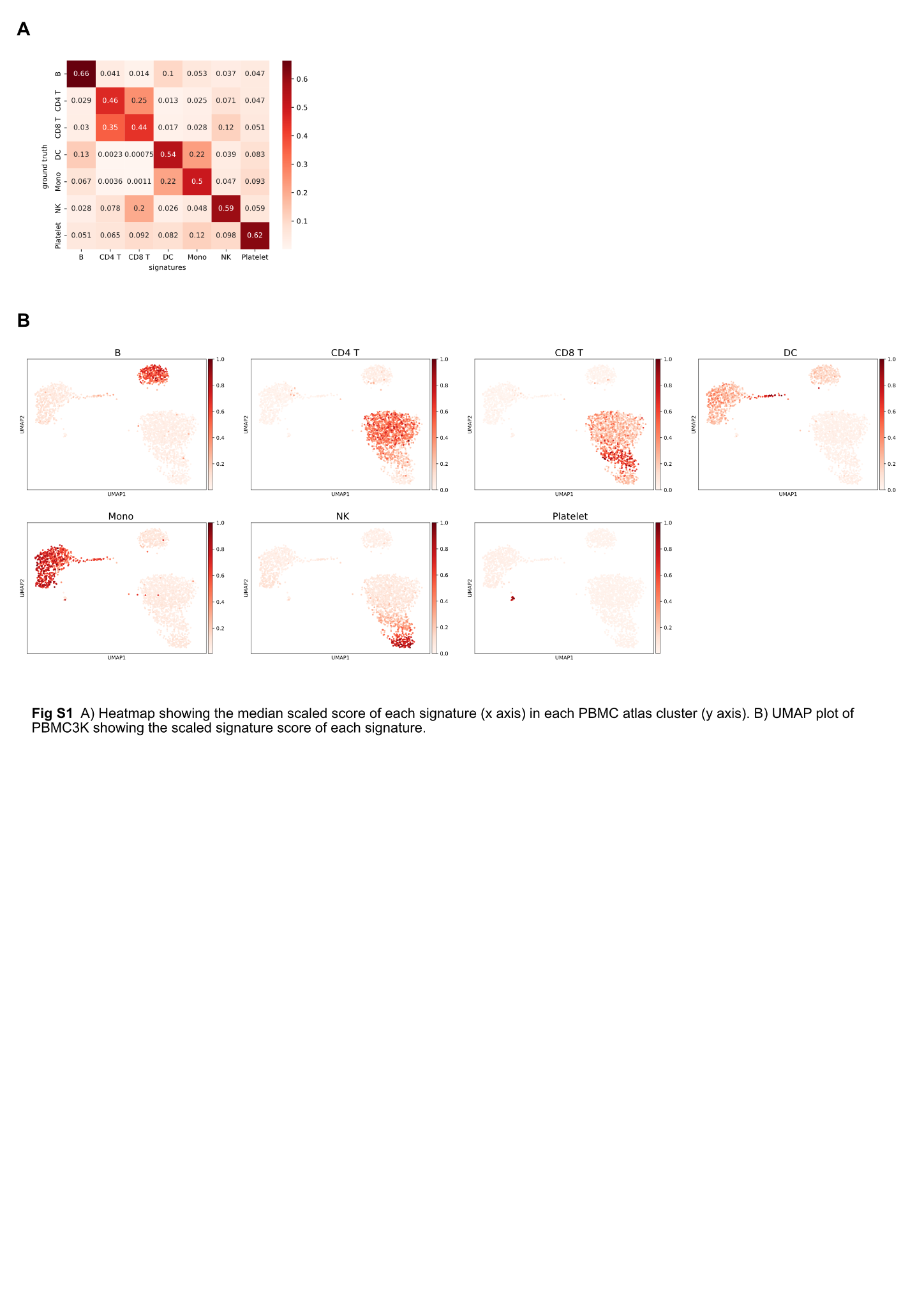

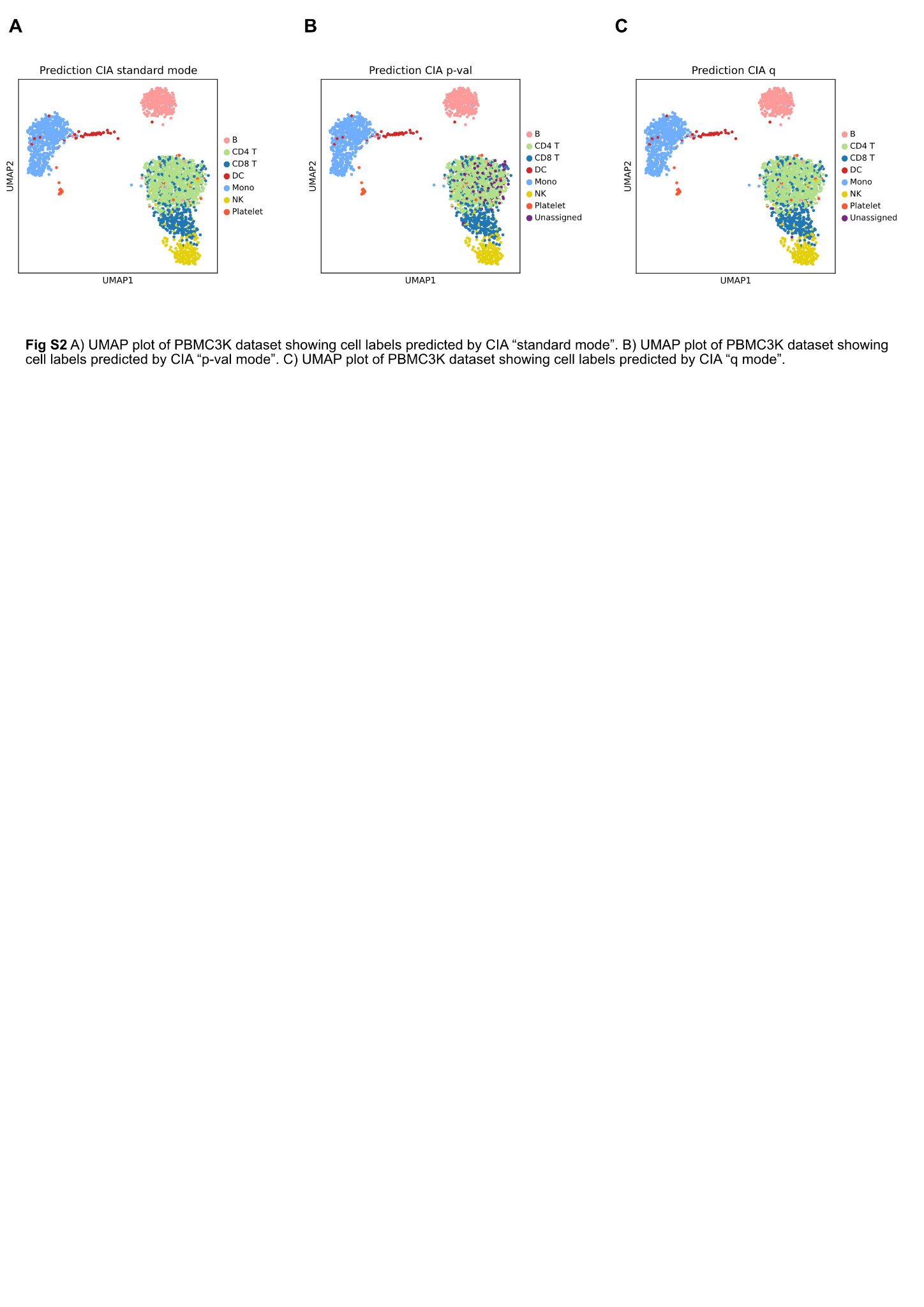
